# Development of a yeast-based assay for bioavailable phosphorus

**DOI:** 10.1101/2021.02.28.433264

**Authors:** Heather A.M. Shepherd, Matt T. Trentman, Jennifer L. Tank, Jennifer Praner, Anissa Cervantes, Priya Chaudhary, Jonah Gezelter, Allyson J. Marrs, Kathryn A. Myers, Jonathan R. Welsh, Yueh-Fu O. Wu, Holly V. Goodson

## Abstract

Preventing eutrophication of inland freshwater ecosystems requires quantifying the phosphorus (P) content of the streams and rivers that feed them. Typical methods for measuring P assess soluble reactive P (SRP) or total P (TP) and require expensive analytical techniques that produce hazardous waste. Here we present a novel method for measuring the more relevant bioavailable P (BAP); this assay utilizes the growth of familiar baker’s yeast, avoids production of hazardous waste, and reduces cost relative to measurements of SRP and TP. The yeast BAP (yBAP) assay takes advantage of the observation that yeast density at saturating growth increases linearly with provided P. We show that this relationship can be used to measure P in freshwater in concentration ranges relevant to eutrophication. In addition, we measured yBAP in water containing known amounts of fertilizer and in samples from agricultural waterways. We observed that the majority of yBAP values were between those obtained from standard SRP and TP measurements, demonstrating that the assay is compatible with real-world settings. The cost-effective and nonhazardous nature of the yeast-based assay suggests that it could have utility in a range of settings, offering added insight to identify water systems at risk of eutrophication from excess phosphorus.

## INTRODUCTION

Eutrophication, a form of water pollution caused by excess nutrients, is a concern for inland freshwater ecosystems worldwide.^1, 2^ Surplus nutrient supply can lead to algal blooms, which disrupt ecosystems, often followed by algal senescence which can lead to hypoxic zones that limit biology; these bloom events impact local economies by reducing commercial fishing, water recreation, and waterfront real estate.^3^ In the Great Lakes region, phosphorus (P) pollution from excess fertilizer application on agricultural lands contributes significantly to recurring algal blooms.^4–7^ Moreover, over application of P-based fertilizer is also costly to farmers and represents a waste of a limited resource.^8^ To prevent eutrophication and reduce P waste, accurate P detection methods are needed to pinpoint where excess P is entering waterways. Standard approaches measure either the total P (TP) or soluble reactive P (SRP); SRP typically provides an artificially low estimate of BAP,^9^ while TP often provides an artificially high estimate.^10, 11^ The quantity most relevant to the biological response to eutrophication is the bioavailable P (BAP), i.e., the portion of P accessible to organisms.^12, 13^ Of the standard analyses, SRP is the easiest to perform: it relies on filtered samples in which forms of P that are both soluble and simple (e.g., orthophosphates) react with molybdic acid to produce a measurable color change.^14, 15^ While straightforward, SRP assays fail to quantify the BAP in complex molecules (e.g., DNA) or the particulate fraction (e.g., adhered to minerals in clay or present in partially decomposed organic materials^16, 17^). In contrast, the TP measurement captures complex or sediment-bound P forms by first subjecting the sample to high temperatures in strong acid^18^ before the molybdic acid reaction.^19^ However, through this process, TP assays can quantify P in forms that are not typically bioavailable. As such, BAP is generally expected to fall between these two standard measurements, and through a combination of biological and chemical processes (e.g., enzyme secretion, local acidification), organisms access more P than is detected by SRP.^12^ However, a portion of the total P typically remains unavailable to organisms, either because it resides as large particles or because the P is in a chemical form that is biologically inert.^10^

Other non-ideal aspects of the SRP and TP assays include that they produce hazardous waste, require lab-intensive techniques, and are expensive.^20^ For example, the molybdic acid reaction used in both assays requires heavy metals (e.g., stannous chloride and molybdenum), and the TP assay also requires high-temperature (e.g., boiling) acid, which is dangerous even in laboratory settings. Charges for outsourcing these assays can range from $5-$20/sample for SRP and $10-$30/sample for TP (M. Trentman, pers comm), which can be cost prohibitive for users trying to monitor their local surface water for P pollution or farmers that want to optimize their fertilizer use. Thus, there is a need for an improved P-detection method that is user-friendly, cost-effective, and that can accurately quantify BAP, rather than SRP or TP.

Other BAP assays do exist and can be loosely categorized as chemical, quasi-biological, and biological. Chemical methods for measuring BAP in water provide an approximation of BAP by releasing P through a combination of chemical extractions and/or enzyme treatments, and then measuring it by one of various methods such as molybdate colorimetry (similar to SRP) or NMR.^13, 16, 21–24^ Chemical methods for assessing BAP have some of the same drawbacks as those for measuring SRP and TP. In addition, the ability of these assays to approximate BAP likely varies with the chemical composition of the particular water sample.

A recently developed quasi-biological method for BAP uses qPCR measurements of phosphatase gene expression in bacteria that are added to water samples,^25^ but the approach requires sophisticated equipment not present in many environmental labs. Also, because the bacteria are exposed to the sample for only short periods (2-8 hours), this assay does not account for slower biological or chemical processes that allow organisms to extract P from sediments in turbid environments. As such, this method may underestimate BAP for at least some types of water bodies.

Other biological methods for detecting BAP include assays based on the growth of algae^10, 12, 26^ or bacterioplankton.^27^ While these approaches link BAP directly to the organisms most relevant to eutrophication, algal assays can take multiple weeks, use relatively large volumes of sample water, and require specialized growth environments.^12^ Bacterioplankton-based methods can be run more rapidly, but are difficult to reproduce due to the uncharacterized nature of the cultures used thus far.^27^ In summary, there is room for improved methodology in measuring BAP.

To address this problem, we designed a BAP assay based on the growth of the baker’s yeast *Saccharomyces cerevisiae*; yeast are safe, easy to work with, and resistant to hazards (e.g., viruses, pesticides) common in aquatic environments.^28, 29^ Additionally, yeast grow quickly and are well-adapted for lab use. The resulting yeast BAP (yBAP) assay produces a linear response at P concentrations between 4 and 40 µM P (i.e., between 31 and 1240 µg L^-1^) and can be extended to higher concentrations (∼80µM) with the use of non-linear fitting functions. We tested the yBAP assay using water samples made from fertilizer, as well as water samples from an agricultural stream. We observed that the measured values fell between SRP and TP, indicating that the yBAP assay is compatible with real-world environmental settings. Moreover, the yBAP assay produces no hazardous waste, can be run with minimal equipment, and reduces cost relative to SRP and TP tests, suggesting that it may be a useful addition to the set of P-quantification tools used to combat eutrophication.

## METHODS

### Yeast Bioavailable Phosphorus (yBAP) Assays

We set out to develop a technique and methodology that uses the density to which a yeast population grows to quantify BAP in environmental water samples; as discussed below, the assay is based on the principle that the saturating level of yeast growth is linear with the concentration of a limiting nutrient.^30^ We developed two variants of a yeast based method to estimate yBAP, each with a different experimental scale; one assay is conducted in 50 mL centrifuge tubes and the other in 200 µL 96 well plates (hereafter referred to as the centrifuge tube and microtiter plate assays, respectively). In both assays, the size of the yeast population that results after growing in the presence of test water supplemented with all required nutrients, except P, is assessed by measuring the optical density. The observed yeast density is then compared to a standard curve prepared with potassium phosphate. The centrifuge tube assay was designed to require minimal equipment, with the goal of making it useful for community-based monitoring efforts and for hands-on instruction in schools. To enable low-cost parallel processing of a larger number of samples, we miniaturized our assay for use in a microtiter plate, instead of a rack of centrifuge tubes, thus increasing the testing potential for analytical labs.

#### Centrifuge Tube Assay

For each individual assay, we placed 5 ml of aqueous sample (unknown or a standard, as described below) in a 50 ml “Falcon” centrifuge tube containing 5 ml 2X phosphate-free yeast growth media (described below), 50 µM (final) ampicillin and kanamycin, and 10 mM (final) sodium citrate/citric acid at pH 4.5; antibiotics were included to discourage bacterial growth, buffers were added because yeast require an acidic environment, and the 40 ml of head space in the tube provided oxygen. Baker’s yeast (*Saccharomyces cerevisiae,* described below) was inoculated into these tubes from a phosphorus-depleted stock culture to achieve a final yeast concentration of 0.05 optical density (OD) at 600 nm. We chose this OD because preliminary experiments with less yeast gave inconsistent results, perhaps because of insufficient enzyme secretion from the smaller populations. We closed the caps tightly and incubated the tubes without shaking at 37°C for four days, at which point the culture density had plateaued. We note that we chose 37°C as an incubation temperature, instead of the typical 30°C for yeast, because 37°C incubators are more readily available. To measure the final yeast density, we mixed the cultures by inversion and measured the absorbance at 600 nm on a spectrophotometer (Genesys 10S UV-Vis, blanked against 1x media made with double deionized (dd)H_2_O).^31^ When performing studies of environmental water samples, the blank (different for each sample) was prepared the day of reading by using fresh sample in place of the deionized water to dilute the media; this process accounted for the possible presence of sediment or other particulates in the samples. We generated standard curves in parallel using known amounts of freshly diluted potassium phosphate. For precision measurements, we ran both experimental and standard samples in triplicate. We describe the curve-fitting process below and include the step-by-step protocol in the supplementary material.

#### Microtiter Plate Assay

We miniaturized the centrifuge tube assay by adapting it to 96 well flat-bottom microtiter plates. The media, antibiotics, water, controls, and P standards were all used in the same ratios and to the same final concentrations as in the centrifuge tube assay above but scaled to a 200 µl culture in microtiter wells. To avoid edge-related artifacts during absorbance measurements, we did not use the exterior wells of the plate. Instead, we filled the 36 outer wells with 200 µl of the double deionized water containing ampicillin and kanamycin (the antibiotics may have not been necessary but were added to maintain consistency across the plate). We incubated the plates without shaking at 30°C for 4 days; we used the optimal yeast growth temperature of 30°C under the assumption that labs with plate readers would likely have 30°C incubators. We took OD readings of the plate (including the blanks) at 600 nm on a Synergy HI plate reader at the University of Notre Dame’s Center for Environmental Science and Technology (CEST) set to 30°C; before reading, we shook samples using an orbital formation for 1 min, at 807 cycles per minute, in order to mix and suspend the yeast in each sample well. Please see supplementary material for step-by-step protocol.

#### Calculations

For both assays, we plotted the absorbances against the known P concentrations to generate a standard curve. We found that it was best to prepare a new standard curve with each set of measurements, though under well-controlled circumstances we found that the curves are highly reproducible. To use this dataset to determine the concentrations of unknown samples, we fit the linear portion of the curve to the equation:

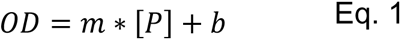

where OD is the optical density of yeast after incubation, [P] is the input phosphorus concentration, and m and b are constants corresponding to the line slope and Y intercept, respectively. To fit the whole growth curve, we used a fractional saturation equation similar to the familiar Michaelis-Menten function of biochemistry^32^

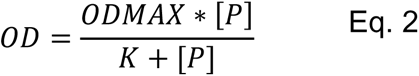

where OD is the optical density of yeast after incubation with a limiting amount of P, OD_max_ is the maximum optical density of yeast after incubation (when P is no longer the limiting nutrient), [P] is the input phosphorus concentration (here analogous to the substrate concentration in an enzyme reaction), and K is the empirically determined half-saturation constant (fitted from the data).

### Yeast Methods

#### Yeast strain

For this work, we utilized *Saccharomyces cerevisiae* strain DBY10148 (a generous gift from David Botstein, Princeton University^30^). We chose this strain because it is prototrophic, meaning that it has metabolic capabilities similar to wild yeast, and flocculates (clumps) less than many other strains.

#### Yeast growth media

For <u>medium-term culture maintenance</u> (weeks-months), we streaked frozen stock cultures of yeast (in 30% glycerol) to single colonies on YPD plates (YPD: yeast-extract peptone dextrose^33^), incubated the plates at 30°C until colonies appeared (a few days), then stored the plates at 4°C indefinitely. For most of our other manipulations in this paper, we used as a foundation <u>phosphorus-free synthetic defined (P_0_SD) media</u> as described previously.^30^ The P_0_SD media was typically prepared as a 2X stock, then diluted to 1x by addition of water (ddH_2_0 or sample water), vitamin/mineral mix, and glucose (1% final) (Table 1). We prepared <u>P-assay media for the yeast assays</u> by supplementing the <u>P_0_SD</u> media with antibiotics and citrate butter (Table 1), which we included in all assays (even those made with sterile lab DI water) to maintain consistency in growth conditions between standard curves and sample testing. To prepare standard curves, we added appropriate amounts of potassium phosphate to this 1x media from a filter-sterilized 1M potassium phosphate solution, which we made freshly every 3 months. For detailed instructions please see the supplemental materials.

**Table 1.** Media composition

| **Media** | **Components for 1L** | **Use** |
| --- | --- | --- |
| **2X Phosphorus-Free Synthetic Defined Base Media (P<sub>0</sub>SD base)** | 0.2 g Calcium<br>0.2 g Sodium chloride<br>1 g Magnesium sulfate<br>0.2 g Potassium chloride<br>10 g Ammonium sulfate | This is the base media for the other two medias. |
| **Assay-Ready P<sub>0</sub>SD Media** | Fill to 1L with 2x P<sub>0</sub>SD base<br>20 g/L Glucose<br>2 ml/L 1000x vitamins*<br>2 ml/L 1000x metals*<br>2 ml/L 1000x ampicillin*<br>2 ml/L 1000x kanamycin*<br>20 ml/L 1M sodium citrate<br>20 ml/L 1M citric acid<br>(concentrations will be halved to 1X with the addition of sample water) | Used in the assays to provide nutrients to the yeast, making P the only limiting factor in the growth. |
| **Phosphorus-Limited SD Media (P<sub>50</sub>SD)** | Fill to 1L with 1x P<sub>0</sub>SD base<br>50 µl 1M potassium phosphate<br>10 g glucose<br>1 ml 1000x vitamins*<br>1 ml 1000x metals* | Used to grow P-depleted stock yeast cultures for assays. |

\*See Supplementary methods for details of the preparation of the vitamins, metals, and antibiotics

#### Preparation of P-depleted Stock Culture

The P-depleted stock culture was grown by inoculating 50 ml of <u>P-limited media</u> (made by supplementing P_0_SD with 50 µM potassium phosphate) with a single colony taken from a typical YPD plate. The culture was then incubated for four days at 30°C, at which time the yeast had reached the stationary phase and were depleted of their internal P stores. These P-depleted cultures were then stored at 4°C for a maximum of two weeks.

### Measurements of P in fertilizer

#### Assays of yBAP in Fertilizer

To test the ability of the assay to handle complex forms of phosphorus, we compared results of both yeast assays between potassium phosphate and pelletized agricultural fertilizer (i.e., phosphorite enriched with phosphorus pentoxide; obtained from Co-Alliance in Buchanan, MI) with a stated N:P:K ratio of 9:23:30. We note that the 23% P in the ratio is in reference to the P_2_O_5_, which is actually 44% P;^34^ this means that the fertilizer is actually 10.1% P by weight. To make a 13.6% (w/v) stock solution, we mixed fertilizer in double deionized water at room temperature for four days with shaking, while covered, to completely dissolve the pellets. These tests were performed as described in the centrifuge tube assay (10 ml cultures) and microtiter plate assay (200 µl cultures) sections above, only replacing the potassium phosphate with appropriately diluted fertilizer solution; we then determined the yBAP content of the fertilizer as described above (see Calculations).

#### Assay of SRP and TP in fertilizer

To compare our yeast assays to an existing analytical method for dissolved P, we measured the soluble reactive phosphorus (SRP) content of the potassium phosphate and the fertilizer using a method adapted from the APHA Standard Methods 23^rd^ edition.^19^ Briefly, we estimated SRP in a 1.5 ml assay using a 2:1:1:1 ratio between water, sample, and 0.2 M molybdic acid, made by dissolving ammonium molybdate in 1.6 M hydrochloric acid (HCl), and 0.1 M stannous chloride (dissolved in 1.4 M HCl), added in the order listed. We took OD measurements at 665 nm after 30 - 45 minutes. To measure total P (TP) in the fertilizer, we sent a fertilizer sample to Brookside Laboratories (New Bremen, OH), where they used method AOAC 2015.18^35^ with Inductively Coupled Plasma–Optical Emission Spectrometry determination (ICP-OES), which corresponds to method “Alternative B” which extracts the TP from the sample by boiling with HCl, before analyzing the sample with ICP-OES.^35^

### Measurements of P in stream samples

#### Sample collection

In order to test the efficacy of the yeast assay on natural water (i.e., not synthesized in the lab), we conducted the yeast assay on samples collected from a nearby agricultural stream (Shatto Ditch Watershed, IN). These samples were collected across the hydrograph of a storm (n=45), and thus likely contain a mix of chemical constituents including organic and inorganic P fertilizers, as well as pesticides and sediments. A total of 11.4 cm of rain occurred during the event, and stream discharge peaked at 1700 L s^-1^; we began sample collection after rainfall started and continued intermittently for a week using an ISCO Automated Water Sampler (3700; Teledyne, Lincoln, NE); we then froze samples until later analysis as described below. A full summary of these samples and the context with which they were collected is available in Trentman et al.;^36^ here we highlight a subset of these data to show the efficacy of the yeast growth assay on samples collected from the natural environment.

#### yBAP Assay on stream samples

We used the miniaturized assays when testing the stream water samples. We created a plate containing the standard curve as described above using environmentally relevant P concentrations from potassium phosphate. We ran the water samples in triplicate as described above. To test for the presence of growth inhibitors that might be present in environmental water samples, we added 20 µM potassium phosphate to each set of samples as an additional control.

#### Comparative measurement of SRP and TP from stream samples

For each stream sample, we also measured the SRP content on filtered samples (Type A/E glass fiber filter; Pall, Ann Arbor, MI; frozen until analysis) using a Lachat Quickchem Analyzer (Hach Company, Loveland, CO) and the molybdate-blue ascorbic acid method (Hach Company, Loveland CO; Method=10-115-01-1-Q). We measured TP on unfiltered samples with an in-line UV persulfate digester also on the Lachat Quickchem Analyzer (Method=10-115-01-3-E).

### Chemicals and supplies

For additional chemicals and supplies, please see the Supplementary Methods.

## RESULTS AND DICUSSION

### Relationship between yeast growth and concentration of potassium phosphate

Our first goal to validate the assay was to establish the relationship between P concentration and yeast growth using known concentrations of P, allowing us to use this relationship to determine the amount of P in unknown samples (Figure 1). The approach is based on previous work by Saldanha et al.,^30^ who studied yeast growth under conditions of limiting nutrients and showed that yeast culture growth at maximum density follows a linear relationship with provided P.

**Figure 1.**
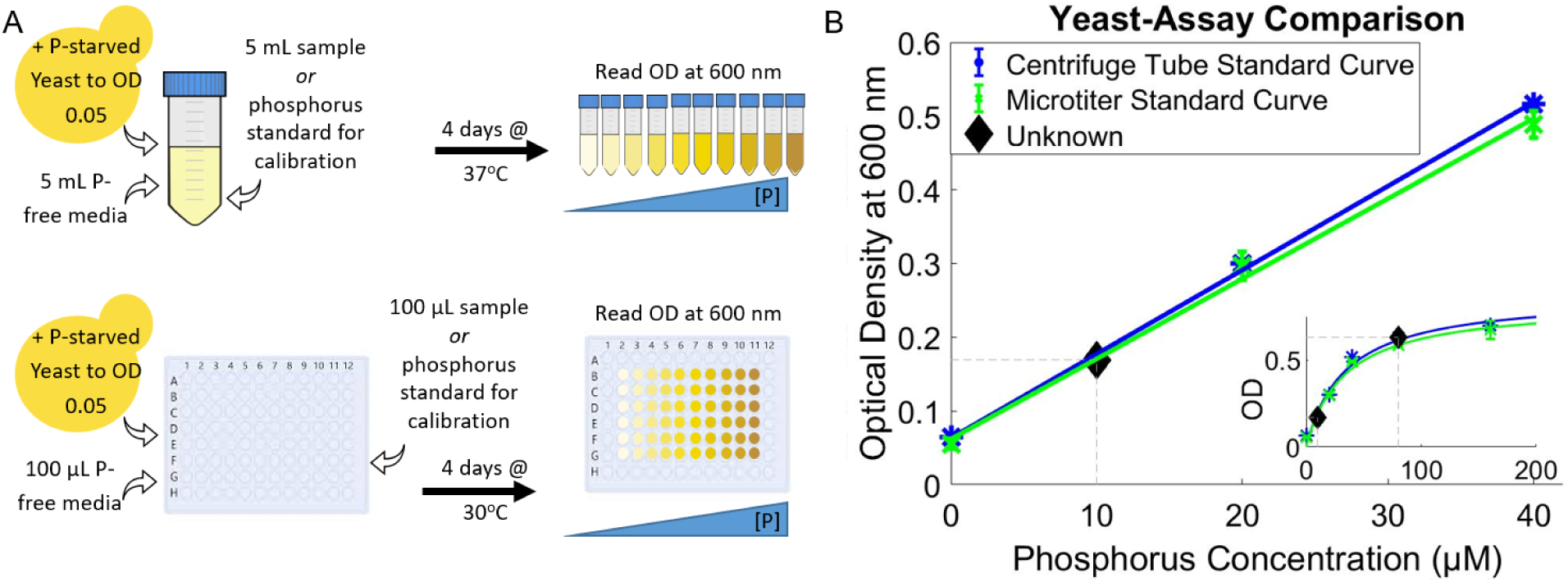
Logic of the yeast-based assay for bioavailable P. (a) Outline of the centrifuge tube and microtiter plate variants of the assay. Briefly, we developed two variants of a yeast-based method to estimate yBAP, each with a different experimental scale; one assay is conducted in 50 mL centrifuge tubes and the other in 200 µL 96 well plates (hereafter referred to as the centrifuge tube and microtiter plate assays, respectively). (b) Using assay data to measure P in unknown samples. Here we show standard curves, constructed for both assay variants using a range of concentrations of potassium phosphate. The standard curve data for the centrifuge tube assay (blue) represent the average +/- S.D. of 5 independent experiments with 3 technical replicates each. The standard curve for the microtiter plate assay (green) represents the average +/- S.D. of 4 independent experiments with 6 technical replicates each. These data show that for both assay variants, yeast growth is linear between 0-40 µM phosphorus (main figure, fitted to y=mx+b) and saturates as P increases (inset, fitted to the fractional saturation function, see Methods). For samples with low P (<40µM), either fit can be used to estimate P from the observed yeast optical density; for samples with higher P, the saturation function is more accurate. To find the P concentration of an unknown sample (hypothetical low and high P examples shown as diamonds), use the appropriate standard curve to determine the P value that corresponds to the optical density observed for that sample. The grey dotted lines associated with each black diamond provide the extrapolation from optical density to the appropriate fit and thus to the measured P concentration. *Notes:* 10 µM = 0.31 ppm= 310 µg L^-1^ P; most error bars on the standard curve datapoints are invisible because they are smaller than the data symbols.

As a first step, we designed a centrifuge tube assay in which yeast were grown to saturation in P-free synthetic defined media with/without known amounts of added potassium phosphate and/or sample water (Figure 1A). We expected the relationship between [P] and yeast density to be linear based on previous work,^30^ which would allow us to determine yBAP via linear extrapolation (Figure 1B). We found the relationship between P and yeast OD as determined from the standard curve to be both linear and highly reproducible for P concentrations between 0 and 40 µM (Figure 1B). This behavior meant that at lower concentrations (<40 µM), the relationship between P and OD could be modeled by a linear fit. However, at higher P concentrations, we began to see saturation behavior (Figure 1B, inset). This saturation occurs because at higher P concentrations, other nutrients eventually become limiting, and at higher yeast concentrations, waste by-products accumulate and inhibit growth.

To extend the useful range of the assay to P concentrations > 40 µM, we fit the data to a fractional saturation equation similar to the Michaelis-Menten equation, a well-known biochemical relationship that describes enzyme saturation.^32^ We observed that this equation fits the data reasonably well (Figure 1B, inset), enabling samples at concentrations above the linear range (e.g., 40-80 µM) to be interpreted by extrapolation from the fitted curve in a manner analogous to the linear extrapolation (Figure 1B).

The initial centrifuge tube-based assay requires limited equipment (an incubator and a spectrophotometer), making it accessible to a wide range of users (e.g., science classrooms). However, it also requires relatively large sample sizes, occupies a large workspace, and produces substantial plastic waste. To address these concerns, we adapted the assay to the wells of a microtiter plate, which gave results very similar to the centrifuge tube assay. We observed the same linear relationship between yeast growth and 0-40 µM potassium phosphate, and again found that the fractional saturation equation fit the data at higher P concentrations (Figure 1B, note green vs. blue curves). These observations indicate that the assays worked as expected with known concentrations of potassium phosphate and thus provided a foundation for attempting to use the yeast-based assay to provide a measure of the concentration of yBAP in experimental samples.

### Relationship between yBAP and input phosphorus in fertilizer

As a first test of the yeast assay to determine P in an unknown sample, we measured the BAP in a sample of P_2_O_5_-based agricultural fertilizer. By comparing the OD of yeast grown with fertilizer to the full standard curve produced with potassium phosphate (Figure 2), we found the yBAP fraction of the fertilizer to be on average 8.7 ± 1.7% using the centrifuge tube assay, and 9.8 ± 1.9% from the microtiter plate assay (Tables S1 & S2). The two methods produced data that were within one standard deviation of each other, supporting the precision of the yeast-based assays.

**Figure 2.**
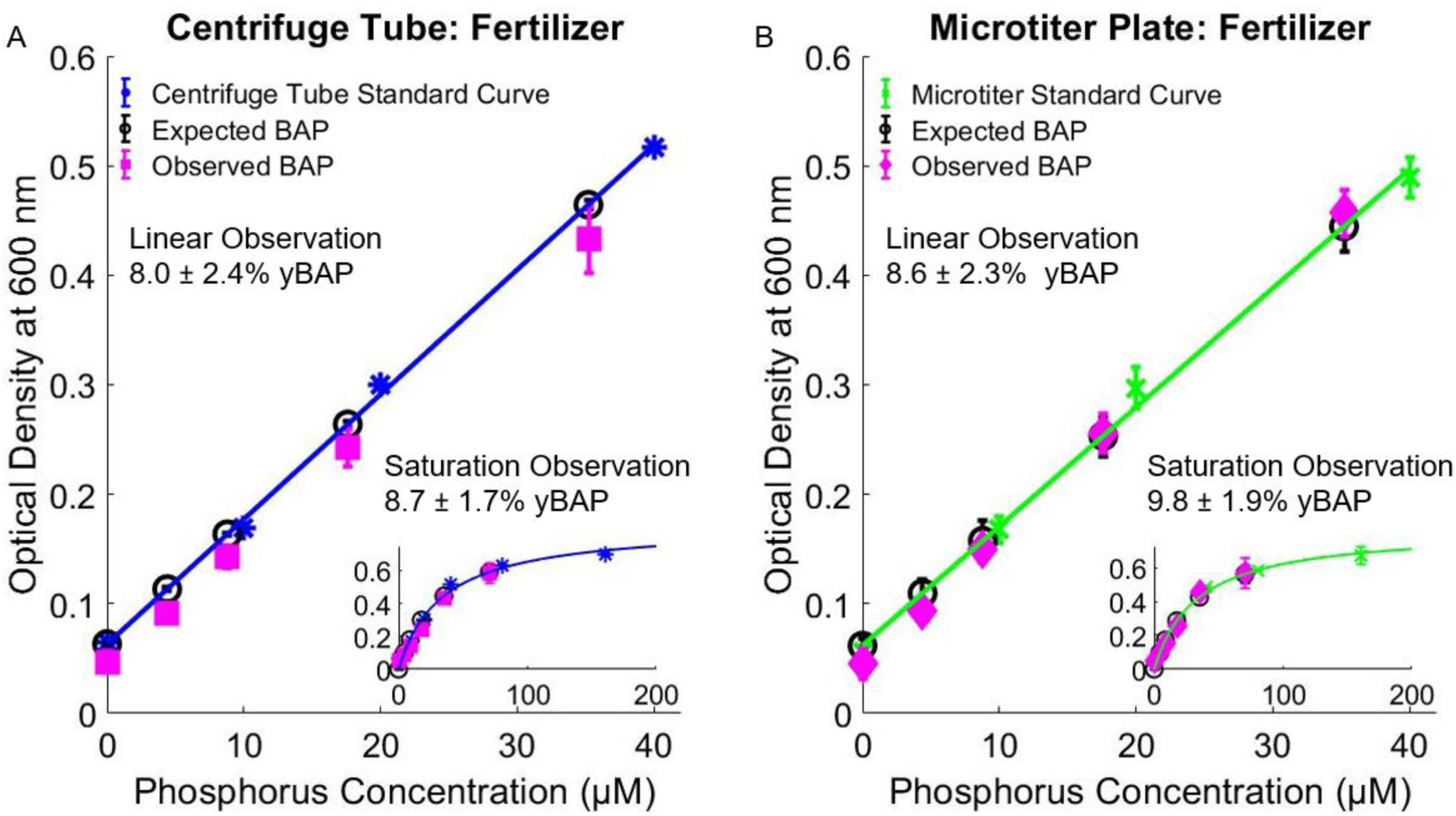
Use of yeast assays to detect phosphorus in agricultural fertilizer. The linear portion of the standard curves is shown in the main graphs (fitted to y=mx+b); the insets are the entire dose response curve (fitted to the fractional saturation function). Error bars provide standard deviation; some are obscured by the data markers. These calibration curves were used to measure the concentration of bioavailable phosphorus in various dilutions of fertilizer as outlined in Table S1. The expected concentrations of the P from the fertilizer were calculated from the 23% P_2_O_5_ (10.1% P), stated by the manufacturer. The data from the various dilutions were averaged together to provide the yBAP measurements indicated above, which are within error of the expected yBAP for both the linear and saturation fits. See Table S1 and Table S2 for individual measurements. Data represent the average of 5 independent experiments with 6 technical replicates each, +/- S.D.

To determine the accuracy of the yeast-based assays, we compared the yBAP values to measurements of SRP and TP in the fertilizer samples, and also the amount of P expected from the documentation provided by the supplier. On average, the SRP content of the fertilizer was 9.7 ± 3.7% P, which was indistinguishable from yBAP. We also analyzed fertilizer TP, which was on average 9.5% total P, again consistent with the yBAP measurement because all of the P_2_O_5_ was expected to be bioavailable. Moreover, all of these measurements were consistent with the P content reported by the manufacturer (10.1% P).

To determine the useful range of the assay, we examined its ability to return the correct value for fertilizer P at different dilutions. We found that both fits to data from both assays worked well in the range between 4 µM and 40 µM P (Figure 2), equating to a range of 124-1240 µg L^-1^. Moreover, we observed that the saturation curve fit to these data provided more accurate measurements in the range between 40 µM and 80 µM P. This range is relevant to environmental P concentrations that promote eutrophication, which are generally described as occurring at concentrations > 3.2 µM (20-100 µg L^-1^ TP^37^). Note that the saturation curve fit could potentially be used to measure even higher P concentrations, but we recommend avoiding trying to interpret OD data that are nearing the plateau of the saturation curve, because at higher P concentrations, the OD changes little with additional input P, creating significant uncertainty in the resulting measurements.

There was general agreement among the yBAP, SRP, and TP measurements for fertilizer, but because we measured both yBAP and SRP for a series of different fertilizer dilutions, we can compare the performance of the two assays at different concentrations. Although the overall differences were small, we observed that the SRP measurement slightly overestimated fertilizer P for concentrations below 16 µM and underestimated fertilizer P for concentrations above 16 µM (Figure 3). In contrast, the yBAP measurements obtained from the fractional saturation curve fit more closely matched the expected value for fertilizer P across the range tested. We note that the yBAP measurements from the linear fit underestimated the 70 µM datapoint (Figure 3), but this is to be expected because the relationship between fertilizer P and yeast density became nonlinear above 40 µM (Figure 1B, inset).

**Figure 3.**
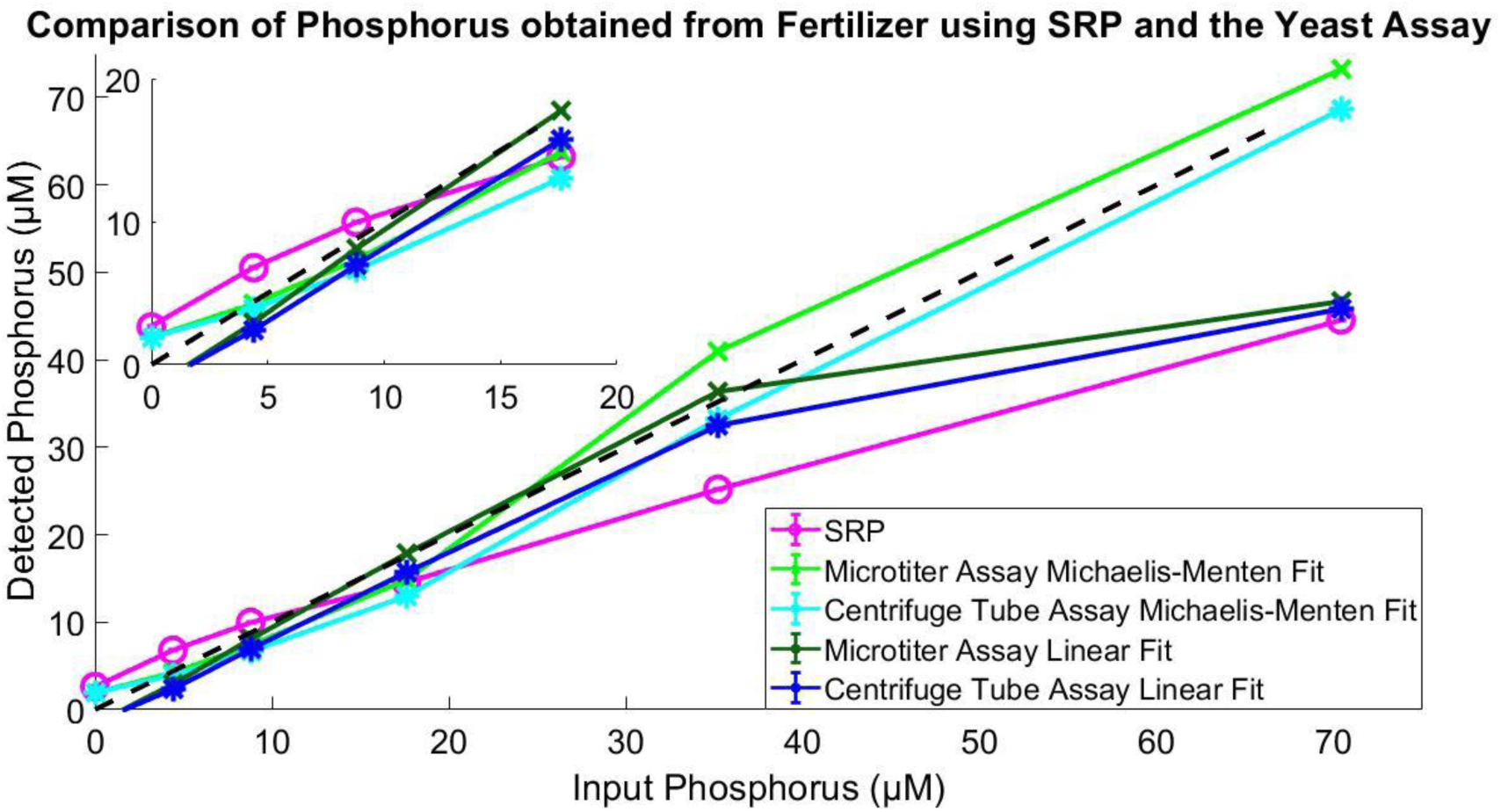
Comparison of fertilizer phosphorus as detected by SRP measurements and the yeast-based assay, using the standard curves in Figure 2. Lines are added to guide the eye. Error bars are standard deviation; most are obscured by the data markers. Input phosphorus (X axis) refers to the concentration of phosphorus in the diluted fertilizer sample, as determined from the manufacturer’s stated P_2_O_5_ content. Detected phosphorus (Y axis) refers to the concentration of P detected by the indicated method. Each datapoint represents the average of 5 independent experiments with 6 technical replicates each. The black dotted line represents data trend expected for a method that could detect all input P. These data show that the SRP can overestimate P for input phosphorus below 16 µM and underestimate P at higher concentrations. In contrast, the fractional saturation fit of the yeast assay was able to more faithfully return P content across the whole range. See Table S1&S2 for additional information related to this plot.

### Detection of yBAP in stream water samples

The yeast-based assays were able to produce measurements of yBAP in fertilizer that were consistent with TP and SRP measurements using standard methods. Next, we tested the utility of our yeast assay with environmental water samples that would potentially have fertilizer P as well as P from other sources. We examined yBAP using n=45 water samples collected from an agricultural stream before, during, and after a storm. These water samples were expected to contain fertilizer run-off from nearby farm fields as well as environmental P from underlying substrate and decaying biomass. Here we compare yBAP using the microtiter plate assay to SRP and TP measurements using standard methods.

We found that all stream water samples fell in the linear range of the standard curve for the microtiter plate (Figure 4) and had BAP concentrations less than 40 µM (<1,240 µg L^-1^). We then compared BAP to the SRP and TP measurements (Figure 4B, C) and found that yBAP concentrations were generally near but slightly higher than SRP, but somewhat lower than TP. Thus the yBAP measurements fell between the TP and SRP measurements (Figure 4), which was consistent with results from measurements made using algal BAP assays.^10^ In contrast, previous bacterioplankton assays have found that BAP was often below both SRP and TP.^27^ Nevertheless, observations from this study provide evidence that the yeast-based assay can be used for added insight on BAP in samples obtained from field settings.

**Figure 4.**
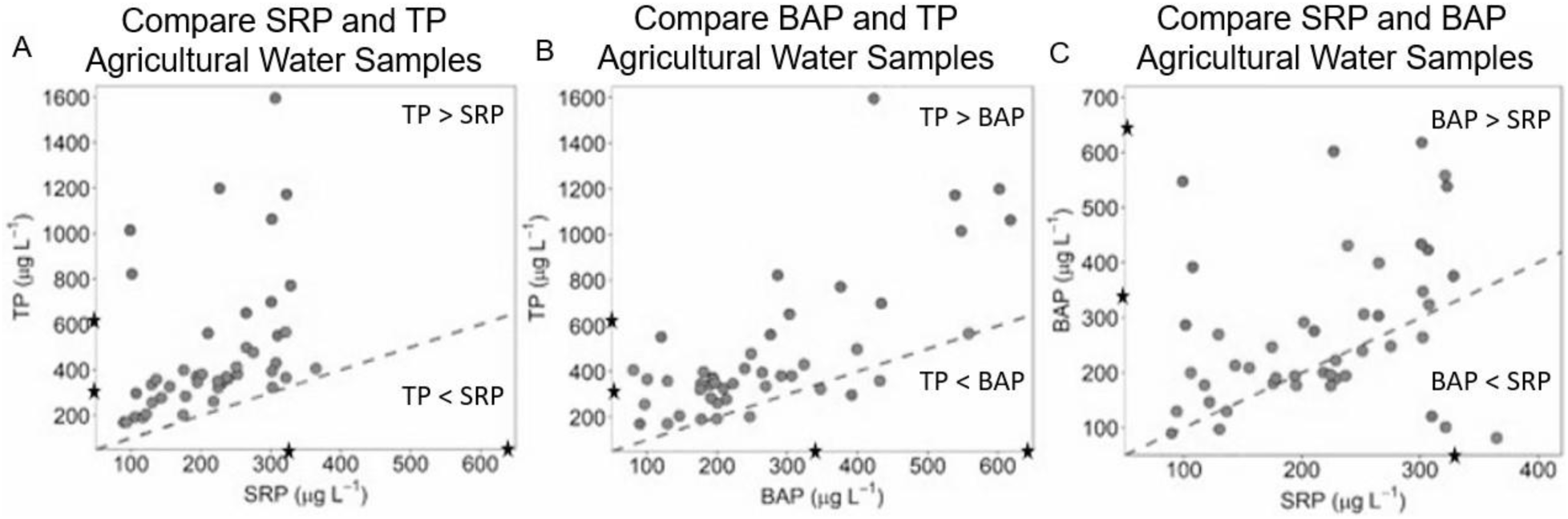
Comparison between methods used to measure P in agricultural water samples. The microtiter plate assay was used to determine the concentration of BAP in agricultural water runoff samples. (a) The TP and SRP of each sample were measured using the Lachat Quickchem Analyzer; these data show that the TP measurements were consistently higher than the SRP measurements. (b) The BAP values, measured using the microtiter plate assay, were consistently lower than TP. (c) The majority of the BAP values were higher than SRP values. Measurements recorded in µg L^-1^. *The stars indicate 310 ug L^-1^ and 620 ug L^-1^, which correspond to 10 µM and 20 µM P respectively.

### Practical recommendations and considerations in using the assay

#### Two variants of the assay

The Centrifuge Tube yBAP Assay was developed to minimize the need for specialized equipment, especially if the prepared culture media is already available (see supplemental methods for materials/supplies). For larger numbers of samples or cases in which the water sample size is limited, the microtiter plate yBAP Assay is preferable, having the same basic supply needs, except for the addition of a plate reader instead of a spectrophotometer.

#### Temperature

As presented here, we grew the centrifuge tube yBAP assay cultures at 37°C (a common incubator temperature used for bacterial cultures) and the microtiter plate cultures at 30°C (the more typical temperature for yeast) but expect that either temperature could be used for either assay.

#### Time

As noted in the methods, we grew the yeast cultures for four days, and we recommend that users not try to shorten the growth time for two reasons. First, the linearity between yeast density and limiting nutrient is established only for the density plateau, and the situation is more complicated if the culture density is still changing when the measurements are taken. Second, it is important to give the culture time to extract P from complex forms and/or sediments. If users do try to adjust the time or other aspects of the conditions, it is important to make sure that the culture densities have reached a plateau before taking the density readings.

#### Possible bacterial contamination

For the reported yBAP assays, we included a combination of antibiotics to reduce bacterial contamination. In our experience, some level of remaining bacterial contamination is common when working with field samples, but small amounts of bacteria do not impact results because bacteria also scatter light, as yeast do. However, we suggest that it is important to include the antibiotics because if a situation arises where most of the light scattering comes from bacteria, rather than yeast, variation caused by presence of environmental bacteriophages (viruses that attack bacteria) could negatively impact the yBAP results.

#### Possible presence of poisons or other yeast growth inhibitors in field samples

As noted above, yeast are known to be resistant to glyphosate-based herbicides.^29^ In addition, they are robust to potential problems such as pH changes as long as the initial starting pH is not basic; we ensured this through the addition of citric acid buffer to the growth media (see Supplemental Methods). While we have demonstrated that the yBAP assay is compatible with field samples from an agricultural stream (Figure 4), it is always possible that a field sample would contain a poison or other growth inhibitor. To control for this problem, we suggest that for each field sample, users include a control sample to which 20 µM potassium phosphate has been added, such that if the amended sample has an OD less than that of the 20 µM standard, then the presence of inhibitors should be suspected.

#### Need for shaking and/or aeration

Yeast are able to grow with very minimal oxygen, such that the yeast does not need aeration during the growth period. However, yeast do settle quickly in suspension, so for reproducible results, it is important to make sure that cultures are well-mixed before reading OD values.

#### Turbidity

Water samples collected from the field often contain sediment or other materials that could interfere with the OD measurement, but we found that the yBAP assay is compatible with moderate amounts of natural turbidity (30-4500 NTU^36^); recall that the 45 field samples were taken during a storm event. During the spectrophotometer readings, we subtracted a turbidity blank based on the water sample from the OD of the yBAP reading. We suspect that as long as the sum of the OD from the yeast and the turbidity is less than ∼0.7 (the point at which most spectrophotometers start to become nonlinear, see Figure S1), low levels of turbidity will not be a problem for measuring moderate levels of yBAP. As a goal for future work, we are planning to develop modified versions of the yBAP assay to be compatible with higher field turbidity through the use of non-optical methods.

## Conclusions

Here we have presented a yeast-based assay for measuring bioavailable P (yBAP) in water samples, and have shown that the range of utility is compatible with the goal of detecting P concentrations relevant to eutrophication of freshwaters (above 3.2 µM P; 20-100 µg L^-1^ TP^37^). When we tested the yBAP assay on water samples collected from an agricultural stream, we found that yBAP concentrations fell between the SRP and TP values measured for the same samples, indicating that the yBAP assay has utility for field samples. Moreover, compared to SRP and TP analyses, the yBAP assay produces less hazardous waste and requires no hazardous laboratory manipulations (e.g., hot acid for TP). The yBAP assay also reduces costs from $5-$30 per sample using the chemical analyses to $2-$12 per sample using the yeast-based methods (see Supplemental Material for cost analysis). Furthermore, by quantifying BAP specifically, our assay adds insight regarding the measurement of the P form that is most relevant to eutrophication. Compared to other bio-assays for BAP, our assay can be run more quickly than those based on algae,^10, 12^ is easier to interpret compared to those based on bacteria,^27^ and requires less sophisticated equipment than qPCR-based approaches.^25^ Taken together, these attributes suggest that the yBAP assay could become a useful addition to the analytical toolbox used to address the eutrophication of freshwaters.

## Acknowledgments

The microplate analyses were conducted at the Center for Environmental Science and Technology (CEST) at the University of Notre Dame, and MTT was supported in part by a CEST Predoctoral Fellowship. This work was supported by the National Science Foundation [NSF DBI-1556349]. We also thank the private landowners for access to the agricultural stream.

## 1. Supplementary Tables and Figures

**Table S1.** Measurements of bioavailable phosphorus in varied dilutions of a commercial fertilizer sample as determined by linear and fractional saturation fits to the centrifuge tube and microtiter plate assays. The conversion of these data to percent P are provided below in Table S2; these data are derived from Figure 2.

| Input Phosphorus based on Manufacturers Label | Centrifuge Tube Assay |  | Microtiter Assay |  | Input Phosphorus based on Brookside TP Data |
| --- | --- | --- | --- | --- | --- |
|  | P from Linear Fit | P from Fractional Saturation Fit | P from Linear Fit | P from Fractional Saturation Fit |  |
| 4.4 $\mu\text{M}$ | 2.4 $\mu\text{M}$ | 3.9 $\mu\text{M}$ | 2.9 $\mu\text{M}$ | 4.2 $\mu\text{M}$ | 4.1 $\mu\text{M}$ |
| 8.8 $\mu\text{M}$ | 7.0 $\mu\text{M}$ | 6.6 $\mu\text{M}$ | 8.1 $\mu\text{M}$ | 7.4 $\mu\text{M}$ | 8.3 $\mu\text{M}$ |
| 17.6 $\mu\text{M}$ | 15.7 $\mu\text{M}$ | 13.0 $\mu\text{M}$ | 17.8 $\mu\text{M}$ | 14.8 $\mu\text{M}$ | 16.6 $\mu\text{M}$ |
| 35.2 $\mu\text{M}$ | 32.4 $\mu\text{M}$ | 33.2 $\mu\text{M}$ | 36.3 $\mu\text{M}$ | 40.9 $\mu\text{M}$ | 33.1 $\mu\text{M}$ |
| 70.4 $\mu\text{M}$ | 45.8 $\mu\text{M}$ | 68.6 $\mu\text{M}$ | 46.7 $\mu\text{M}$ | 73.2 $\mu\text{M}$ | 66.2 $\mu\text{M}$ |

**Table S2.** Calculation of the %P in a commercial fertilizer sample based on the data from varied dilutions of this fertilizer as presented in Table S1.

| Input Phosphorus | Centrifuge Tube Assay |  |  |  |  |  | Microtiter Plate Assay |  |  |  |  |  |
| --- | --- | --- | --- | --- | --- | --- | --- | --- | --- | --- | --- | --- |
|  | Linear |  |  | Fractional Saturation |  |  | Linear |  |  | Fractional Saturation |  |  |
|  | Average | Lower Bound | Upper Bound | Average | Lower Bound | Upper Bound | Average | Lower Bound | Upper Bound | Average | Lower Bound | Upper Bound |
| 4.4 $\mu\text{M}$ | 5.6% | -0.2% | 12.7% | 9.0% | 3.9% | 11.9% | 6.1% | -0.3% | 13.3% | 9.7% | 4.0% | 12.4% |
| 8.8 $\mu\text{M}$ | 8.1% | 4.7% | 12.3% | 7.6% | 3.3% | 10.0% | 8.7% | 5.3% | 13.0% | 8.5% | 3.5% | 10.5% |
| 17.6 $\mu\text{M}$ | 9.1% | 6.9% | 11.7% | 7.5% | 3.5% | 9.4% | 9.7% | 7.4% | 12.5% | 8.5% | 3.8% | 10.1% |
| 35.2 $\mu\text{M}$ | 9.3% | 7.7% | 11.3% | 9.6% | 5.8% | 10.8% | 9.9% | 8.3% | 11.9% | 11.8% | 6.8% | 11.9% |
| 70.4 $\mu\text{M}$* | 6.6% | 5.6% | 7.8% | 9.8% | 9.5% | 11.9% | 6.4% | 5.4% | 7.58% | 10.5% | 9.1% | 9.6% |
| **Average** | **8.0%** | **4.8%** | **12%** | **8.7%** | **5.2%** | **10.8%** | **8.6%** | **5.2%** | **12.7%** | **9.8%** | **5.4%** | **10.9%** |

\*The 70.4 $\mu\text{M}$ input P measurements taken using the linear fits were not included in the average percentages for those fits because 70.4 $\mu\text{M}$ is outside of the linear range.

**Figure S1.**
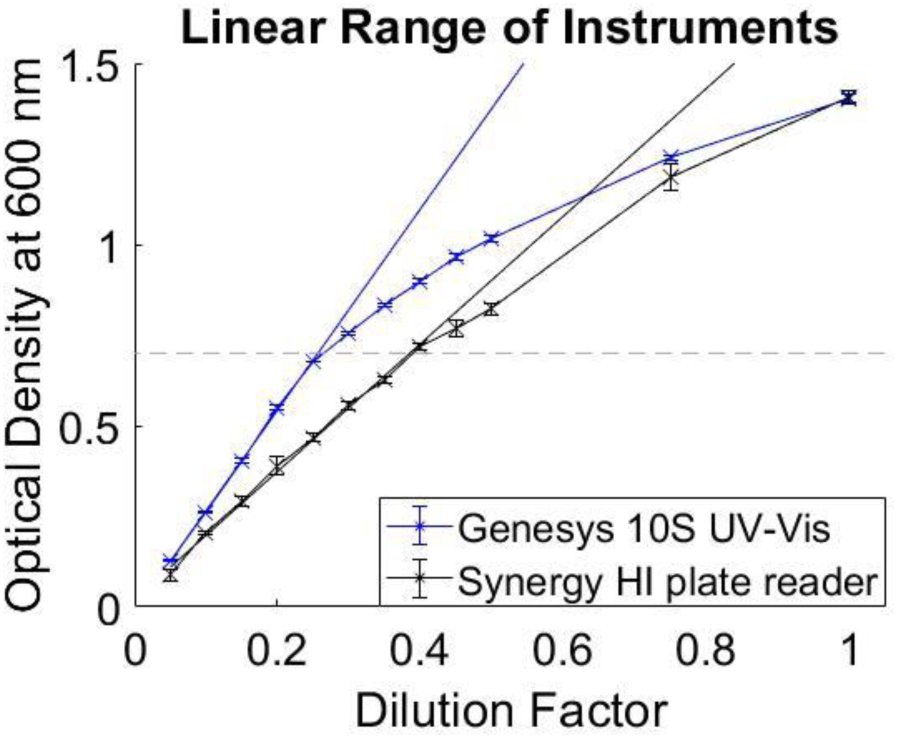
A single yeast culture was diluted to test the linear range of each instrument. The yeast culture was diluted as indicated, and the OD600 was recorded on the specified instrument. These data show that both the UV-Vis and the Synergy HI plate reader loose linearity near OD = 0.7, indicated by the gray dotted line. n=3 or 6 for the centrifuge tube and microtiter plate samples respectively; error bars provide the standard deviation; some are obscured by the symbols.

## 2. Supplementary Methods

### 2.1. Sources of Chemicals and Materials

#### Media Components

*Phosphorus-Free Media Base components:* Sodium chloride (Cat# vw6430-7) and glucose (Cat# 0188) were purchased from VWR International. Potassium chloride (Cat# P217) was purchased from Fisher Scientific (Waltham, MA). Calcium chloride (Cat# 223506), magnesium chloride (Cat# 8.14733.0100), ammonium sulfate (Cat# A7523) were obtained from Sigma Aldrich (St. Louis, MO).

*Vitamins:* Inositol (Cat# A13586), riboflavin (Cat# A11764), and thiamine hydrochloride (Cat# A19560) were purchased from Alfa Aesar (Haverhill, MA). Biotin (Cat# B4501), calcium pantothenate (Cat #CS731), folic acid (Cat# F7876), niacin (Cat #N0761), p-aminobenzoic acid (Cat #100536), and pyridoxine hydrochloride (Cat #P6280) were obtained from Sigma Aldrich.

*Metals:* Boric acid (Cat# BP168-1), potassium iodide (Cat# P410), and magnesium sulfate (Cat# M65) were purchased from Fisher Scientific. Copper sulfate (Cat# 209198), ferric chloride (Cat# 236489), and sodium molybdate (Cat# M1003) were purchased from Sigma Aldrich.

*Antibiotics:* Ampicillin (Cat# AAJ60977-06) was purchased from VWR. Kanamycin sulfate (Cat# BP906-5) was purchased from Fischer Scientific.

*Buffers*: Sodium citrate (Cat# S279) was purchased from Fischer Scientific. Citric acid (Cat# 0529) was purchased from VWR International (Radnor, PA).

*SRP Components*: Ammonium molybdate tetrahydrate (Cat# M1019) and stannous chloride (Cat# 208256) were purchased from Sigma Aldrich.

#### Materials

Centrifuge tubes (Cat# 21008-951) and disposable cuvettes (Cat# 97000-586) were purchased from VWR International. Costar 3596 (Cat# EF5353F-CS), 96-well clear flat-bottom plates were purchased from Daigger Scientific (Buffalo Grove, IL).

### 2.2. Preparation of Growth Media

*Note: If members of the community would like to try our assay but are inhibited by difficulties acquiring or mixing these materials, we can provide aliquots of the metals and vitamins as well as samples of media*.

**Preparation of Phosphorus-Free Synthetic Defined Media (P_0_SD) (1X)** (Saldanha *et al.,* 2004).

The media was typically prepared to a 2X stock and diluted to 1X media by sample water. To prepare the 2X stock please double all ingredients below (see Table 1).

Fill a 1 L graduated cylinder to 500 ml with double deionized water (ddH_2_O), add 0.1 g calcium chloride, 0.1 g sodium chloride, 0.5 g magnesium chloride, 0.1 g potassium chloride and 5 g ammonium sulfate. Fill to 1 L with ddH_2_O. Aliquot 500 mL into 2-1 L borosilicate glass bottles. Autoclave for 15 minutes at 15 psi, 121°C. Store at room temperature in the dark.

#### Preparation of Glucose

A stock solution of 20 % glucose is made by dissolving 10 g of glucose powder in 50 ml of deionized water, add start stir bar before adding liquid; stir powder in slowly to avoid clumps. Filter sterilize the solution using a 0.22 µm filter, and store at room temperature in the dark. When ready to use media, add 25 ml of 20% glucose to 500 ml media for a final concentration of 0.95% glucose; normally [glucose] is 2% in synthetic defined media, but we are following Saldanha *et al.,* 2004.

#### Preparation of Metals

Metals were made to 1000X concentration: Fill a 500 mL graduated cylinder to 250 ml with ddH_2_O, add 500 mg boric acid, 40 mg copper sulfate, 100 mg potassium iodide, 200 mg ferric chloride, 400 mg magnesium sulfate, and 200 mg sodium molybdate. Fill to 500 mL with ddH_2_O. Transfer to a 1 L borosilicate glass bottle. Autoclave for 15 minutes at 15 psi, 121°C. Store at room temperature in the dark.

#### Preparation of Vitamins

Vitamins were made to 1000X concentration:

Fill a 500 mL graduated cylinder to 250 ml with ddH_2_O, add 1 mg biotin, 200 mg calcium pantothenate,1 mg folic acid, 1000 mg Inositol, 200 mg Niacin, 100 mg P-aminobenzoic acid, 200 mg pyridoxine hydrochloride, 100 mg riboflavin, and 200 mg thiamine hydrochloride. Fill to 500 mL with ddH_2_O. Transfer to a 1 L borosilicate glass bottle. Autoclave for 15 minutes at 15 psi, 121°C. Store at room temperature in the dark. Once the above solutions are prepared practice sterile technique during use by using only sterile pipettes to transfer media, flaming the mouth of the bottles before and after pouring liquid and working around a flame.

**Preparation of Assay-Ready Media** - typically made 50 ml at a time, but can be adjusted for the number of samples being tested.

Combine 100 µl 1000x vitamins and metals, 100 µl 1000x ampicillin and kanamycin, 1 ml 1 M sodium citrate, 1 ml 1 M citric acid, and 5 ml 20% glucose in a 50 ml graduated cylinder. Fill to 50 ml with the 2X phosphorus-free base media described above. Note these proportions will be appropriately diluted to 1x by addition of samples or calibration water in the assay.

#### Preparation of Phosphorus-Limited Media

To bring phosphate levels up for growing stock cultures, phosphate-limited media was made by adding glucose, vitamins, and minerals to the diluted 1X phosphorus-free base media as described above, and then supplementing with freshly diluted potassium phosphate to a final concentration of 50 µM. Preparation of samples for standard curves is performed in a similar way. Note that any dilutions of 1M potassium phosphate should be made fresh each day.

### 2.3. Preparation of Phosphorus-Depleted yeast cultures

Inoculate 50 ml of 1X phosphorus-limited SD media with a single colony taken from a typical YPD plate and incubate for four days at 30°C, at which time the yeast have reached the stationary phase. Allowing the yeast to reach stationary phase depletes their internal P storage so that their only P source will be from the sample or standard used in the assay.

### 2.4. Cost Analysis

#### Microtiter plate assay

- $20/hr. technician time; ∼1.5 hours to complete a plate
- $5.68 per microtiter plate
- ∼$2 in assay-ready media
- **Total: $37.68 per full plate** of 60 usable wells **(6 standard curve wells and 18 sample tests,** each paired with two controls; 54 wells assumes outer 36 wells are left empty to avoid edge-related artifacts)
- **$2.10 per sample**. The cost could potentially be further reduced if more samples were run per plate, e.g., by using the outer wells of the plate. Another possibility would be to leave out the positive controls where known [P] is added to the sample to control for the presence of possible growth inhibitors. However, unless users are confident in the absence of any possible inhibitors, we do not advise this.

#### Centrifuge tube assay

- $20/hr. technician time; ∼2 hours to complete a standard set as done in our lab (24 tubes, corresponding to 6 standard curve tubes, plus six samples, each paired with two control tubes)
- $0.81 per tube = $19.39
- ∼$12.50 in assay-ready media for a set of 24 tubes
- **Total: $71.89 for a set of 24 tubes, with six samples**
- **$11.98 per sample.** As above, the cost could potentially be reduced if more samples were run per set, e.g., by leaving out the positive controls where known [P] is added to the sample to control for the presence of possible growth inhibitors. However, unless users are confident in the absence of any possible inhibitors, we do not advise this.

### 2.5. General Notes to be considered before performing either assay

- Practice good pipetting, no bubbles.

○ The presence of bubbles in pipetting alters the true volume being added; this can result in incorrect proportions of media and samples and cause aberrant results.
- Dispose of tips after each use to avoid cross-contamination.
- If using a multichannel pipette with the microtiter plate, make sure each tip is secured and that they all intake the same volume of solution.
- Always agitate yeast mixture before aliquoting to different tubes or wells.

○ Yeast settle quickly. Proper mixing will ensure the amount of yeast delivered to each assay is the same. Failure to mix can result in aberrant results.
- Media should be stored in glassware that has been washed using phosphate-free soap and acid rinsed 3 times using 0.5 M HCl to remove any residual phosphorus.
- Contamination (e.g., from P-containing soap or remnant media) can cause considerable interference in measurements, Thus, we recommend that plastic materials be used one time only to prevent cross-contamination and residual phosphorus carry-over between experiments. If users do attempt to reuse plastic, it is essential that it be washed thoroughly.
- Laboratory yeast cultures can be poured down the drain after being sterilized by addition of bleach or alternatively by autoclaving.

### 2.6. Centrifuge Tube Assay

#### Equipment needed

- Visible light spectrometer
- 1000 µl pipetteman
- 100 or 200 µl pipetteman
- Serological pipette bulb

#### Supplies needed

- 3 centrifuge tubes (50 ml) for each sample (one for the sample and two for sample-specific controls) being tested (note: 15 ml centrifuge tubes will not work as they do not provide enough room for respiration)
- 1.5 ml plastic cuvettes for reading optical density
- 5 ml serological pipettes
- 50 ml serological pipettes
- 1000 µl pipette tips
- 100 or 200 µl pipette tips

#### Control Samples

1. Phosphorus-free media without yeast but with test water (or diluted to 1X with ddH_2_O^1^ for standard curves)

a. This control (one for each sample) is used as a background blank, the value of which is subtracted from the OD reading for that sample<u>. It is assembled immediately before plate reading</u> to avoid the possible effects of microbial growth in the sample.
2. Media with yeast and 5mls of ddH_2_O containing 20 µM P (diluted from freshly made 1 mM potassium phosphate)

a. Comparison of this control (one for each sample) to the standard curve sample with 20 µM P will enable the user to determine whether possible growth inhibitors are present in the test sample.
3. Media with yeast and no phosphorus or test water (media diluted by ddH_2_O instead of test water)

a. This control (one per set of tubes) will indicate the presence of phosphorus contamination in media or glassware. If the yeast remained at the starting OD, no contamination was observed. Note that this tube will also serve as the zero P point for the standard curve.

#### Assay Samples

1. Prepare 5 ml of assay-ready media for each tube being used, but do not distribute it yet.

a. To perform a set of 24 tubes (6 standards, and 6 samples, with two sample-specific controls each) would require 120 ml of assay-ready media total. However only 90 ml of media (18 tubes worth) are needed to start the assay, since the negative controls (sample blanks) will be made the day of the final reading to avoid possible interference by bacterial growth.
2. Add phosphorus-depleted yeast to the prepared assay-ready media; use an amount sufficient to achieve a final optical density of 0.05 at 600nm.

a. For the 24 tubes described above, 18 require yeast (the blanks do not). Thus, for a starting yeast concentration of 0.650, you would add 6.92 ml of yeast to the 90 ml of assay-ready media.
3. Prepare 5 ml of each P calibration standard in ddH_2_O.

a. A typical standard curve would include 5 µM, 10 µM, 20 µM, 40 µM, and 80 µM P. Note that after this 5 ml is mixed with media, the final [P] will be halved, but because the samples will be similarly diluted, this does not cause problems.
b. P should be diluted from 1 mM potassium phosphate freshly made that day from 1M potassium phosphate.
4. Aliquot 5 ml of yeast and media into each 50 ml centrifuge tube (except those being saved for the sample blanks)
5. Add 5 ml of control water, calibration standard or water sample to each tube containing media and yeast.
6. Incubate at 37°C without shaking for 4 days.

a. Though lack of shaking and 37° are unusual for yeast, we designed the assay to be done at 37° and without shaking to minimize equipment needs.
b. We do not recommend shortening the time of incubation because the yeast may need time to extract P from complex substrates.
c. *After 4 days of incubation:*
7. For each sample, prepare a blank by filling a tube with 5 ml of assay-ready media and 5 ml of sample (plus one with ddH_2_O). This should be done immediately before the optical density is taken.
8. Read optical density of all tubes at 600 nm.

a. Blank each sample using the mixture of freshly diluted assay-ready media and the test water mentioned above.
b. Mix each sample immediately before reading to prevent yeast settling.

### 2.7. Microtiter Plate Assay

#### Equipment needed

- Plate reader with 600 nm filter
- 1000 µl pipetteman
- 100 or 200 µl pipetteman
- Serological pipette bulb

#### Supplies needed

- 1 – 96 well flat bottom plate per 18 samples (200 µl wells)
- 5 ml serological pipettes
- 50 ml serological pipettes
- 1000 µl pipette tips
- 100 or 200 µl pipette tips

#### Control Samples

1. Phosphorus-free media (diluted to 1X by test water or ddH_2_O^2^ for standard curves)

a. This control (one for each sample) is used as a background blank, the value of which is subtracted from the OD reading for that sample. It is assembled immediately before plate reading to avoid the possible effects of microbial growth in samples.
2. Media with yeast and 100 µl ddH_2_O containing 20 µM P (diluted from freshly made 1mM potassium phosphate)

a. Comparison of this control (one for each sample) to the standard curve sample with 20 µM P will enable the user to determine whether possible growth inhibitors are present in the test sample.
3. Media with yeast and no phosphorus or test water (media diluted by ddH_2_O instead of test water)

a. This control (one per plate) will indicate the presence of phosphorus contamination in media or glassware. If the yeast remained at the starting OD, no contamination was observed. Note that this well will also serve as the zero P point for the standard curve.
4. ddH_2_O containing 100 µM ampicillin and kanamycin is used to fill the outer wells.

a. This controls for potential edge-related artifacts during absorbance measurements. We did not use the exterior of the plate. Instead, we filled outer wells with 200 µl of the double deionized water containing 100 µM ampicillin and kanamycin (the antibiotics may not be necessary but are added to maintain consistency across the plate).

#### Assay Samples

1. Prepare 100 µl of assay-ready media for each well being used, but do not distribute it yet.

a. To perform a full plate (6 standards, and 18 samples (with two sample-specific controls each)), would require 6.0 ml of assay-ready media total. However only 4.2 ml of media is needed to start the assay, as the negative controls (sample blanks) will be made the day of the final reading to avoid possible interference by bacterial growth.
2. Add phosphorus-depleted yeast to the prepared assay-ready media; use an amount sufficient to achieve a final optical density of 0.05 at 600 nm.

a. For the plate described above, 42 wells require yeast (the blanks do not). Thus, for a starting yeast concentration of 0.650, you would add 323 µl of yeast to the 4.2 ml of assay-ready media.
3. Prepare 100 µl of each P calibration standard in ddH_2_O.

a. A typical standard curve would include 5 µM, 10 µM, 20 µM, 40 µM, and 80 µM P. Note that after this 100 µl is mixed with media, the final [P] will be halved, but because the samples will be similarly diluted, this does not cause problems.
b. P should be diluted from 1 mM potassium phosphate freshly made that day from 1 M potassium phosphate.

i. We suggest making no less than 50 ml 1 mM P to avoid errors in the standards related to pipetting smaller amounts for this dilution.
4. Aliquot 100 µl of yeast and media into microtiter plate wells (except those being saved for the sample blanks).
5. Add 100 µl of control water, calibration standard or water sample to each well containing media and yeast.
6. Add 200 µl of ddH_2_O containing 100 µM antibiotics to outer wells.
7. Incubate at 30°C without shaking for 4 days.
8. For each sample, prepare a blank by filling a well with 100 µl of assay-ready media and 100 µl sample (plus one with ddH_2_O). This should be done immediately before the optical density is taken.
9. Read optical density of the plate at 600 nm.

a. Blank each sample using the mixture of freshly diluted assay-ready media and the test water mentioned above.
b. Mix each sample immediately before reading to prevent yeast settling.

## Footnotes

1 We use double-deioinzed water, but double-distilled or any suitable lab water should work

2 We use double-deioinzed water, but double-distilled or any suitable lab water should work

